# Highly efficient cardiac differentiation and maintenance by thrombin-coagulated fibrin hydrogels enriched with decellularized porcine heart extracellular matrix

**DOI:** 10.1101/2020.01.30.927319

**Authors:** Fatemeh Navaee, Philippe Renaud, Thomas Braschler

**Affiliations:** Laboratory of Microsystems 4, STI-IMT, Station 17, EPFL, 1015 Lausanne, Switzerland; Department of Pathology and Immunology, Faculty of Medicine, University of Geneva, CMU, Rue Michel-Servet 1, 1211 Genève 4, Switzerland

**Keywords:** dECM-fibrin hydrogel, neonatal cardiomyocyte culture, H9c2 cell differentiation, beating synchrony, 3D co-culture

## Abstract

We provide a blend of cardiac decellularized extracellular matrix (dECM) from porcine ventricular tissue and fibrinogen for the formation of an *in-vitro* 3D cardiac cell culture model. Rapid and specific coagulation with thrombin allows gentle inclusion of cells while avoiding sedimentation during formation of the dECM-fibrin composite. We use the system in co-culture with Nor-10 fibroblasts to enhance cardiogenic differentiation of the H9c2 myoblast cell line. The combination of co-culture and appropriate substrate allows to abrogate the use of retinoids, classically considered necessary for cardiogenic H9c2 differentiation. Further enhancement of differentiation efficiency is obtained by 3D embedding. We then proceed with culture of rat neonatal cardiomyocytes in the 3D system. While for H9c2 cells, the collagen content of the dECM was the key factor required for efficient differentiation, the use of dECM-fibrin has specific advantages regarding the culture of neonatal cardiomyocytes. Calcium imaging and analysis of beating motion both indicate that the dECM-fibrin composite significantly enhances recovery, frequency, synchrony and maintenance of spontaneous beating, as compared to various controls including matrigel, pure fibrin and collagen I, but also a fibrin-collagen I blend.

## 1. Introduction

Cardiovascular disease, and in particular myocardial infarction, is the first cause of mortality by representing 31% of global death[1]. In myocardial infarction, cardiac ischemia leads to oxygen deficiency, cell injury and ultimately cell death[2]. After myocardial infarction, structural and functional remodeling in the remaining heart tissues occurs[3]. As adult cardiomyocytes have limited regeneration capability, finding strategies to enhance heart tissue regeneration can help patients recover after myocardial infarction[4]. Our aim here is to provide a simple, yet highly efficient *in-vitro* model for cardiomyocyte differentiation and maintenance. We design this model to simultaneously recapitulate three main dimensions of the native cardiomyocyte environment[5][6]: extracellular matrix cues[7], mechanical stiffness[8] and interaction with stromal fibroblasts[9]. This model has potential applications in drug screening, simplifies generation of cardiomyocytes and may provide building blocks for cell transplantation in regenerative medicine.

Our first cardiomyocyte environment element is the choice of a specific cardiac extracellular matrix. We opt here for the use of porcine decellularized extracellular matrix (dECM) of cardiac origin to provide organ-specific cues[10], with reinforcement by thrombin-coagulated fibrin[11] for improvement of mechanical properties[12].

Myocardial dECM has indeed been shown to powerfully support cardiomyocyte differentiation and maturation in many cell lineages, including human embryonic stem cells and rat neonatal ventricular cardiomyocytes[7]. Moreover, porcine cardiac dECM hydrogels are ultrastucturally similar to their human analogs[13] and known to provide functional benefits after myocardial infarction in animal models[14]. Preservation of part of the organ-specific cues[15][16], [17], including a fraction of the growth factors contained in the dECM[7][18], is thought to be responsible for correct organ-specific cellular differentiation.

Besides dECM, a variety of synthetic and natural matrices have been used for cardiomyocyte culture[19][20][21][22][23]. Collagen I, and mixtures of collagen I with matrigel have in particular been reported to be supportive of spontaneous synchronized *in-vitro* contractions, albeit requiring mixed cardiac cultures with both contractile and non-contractile cells, either primary or derived from induced pluripotent stem cells[20][24][23]. *In-vivo*, collagen foams and matrigel were further shown to enhance engraftment efficiency of the cardiac model cell line H9c2[25]. With regard to the extracellular matrix element, an important aim of this study is therefore to quantify possible advantages of the proposed dECM-fibrin composite over readily available commercial controls such as collagen I or matrigel.

The second element of our *in-vitro* cardiomyocyte niche model is the achievement of appropriate mechanical stiffness. Pure cardiac dECM hydrogels have long gelation times[18] and mechanical properties far below native cardiac tissue[26][7]. This is a practical problem, since sedimentation during the extended gelling time will make cell seeding difficult[27]. More importantly, however, it is known that the differentiated phenotype of heart cells is the most prominent on substrates with stiffness comparable to that of the native heart[8].

Various approaches to reinforce cardiac dECM are available, and include for instance covalent crosslinking[28] or combination with additional hydrogels[29]. Williams, *et al*. combined the two approaches by blending fibrin with dECM from neonatal and adult rat hearts, followed by crosslinking with transglutaminase[30]. However, the lack of specificity of transglutaminases rises concerns about cellular toxicity *in-vitro*[30] and possible side effects such as autoimmune reaction due to altered self-epitopes *in-vivo*[31]. To avoid such side reactions, we here use fibrin obtained by coagulation of fibrinogen with human thrombin[11] to raise the level of stiffness in dECM-fibrin blends in a more specific and safer way. The use of fibrin as opposed to synthetic polymers[29] is motivated by the reported increased efficiency of cardiac reprogramming in the presence of this biopolymer[32].

Finally, the third element of the cardiomyocyte niche to be addressed in this study is the stromal support by fibroblasts. The heart indeed consists only to some 30% of contractile cardiomyocytes[9], the remainder being mainly endothelial cells and fibroblasts[33]. The fibroblast cells in native heart tissue have a major and complex roles in cardiac development, myocardial structure, cell signaling, and electromechanical function[9][34]. *In-vitro*, co-culture with fibroblasts has been shown to enhance skeletal muscle regeneration from myogenic progenitor cells[35] as well as electrophysiological maturation of iPSC-derived cardiomyocytes [36]. We hypothesize here that co-culture with fibroblasts in the context of a mechanically and biologically relevant matrix could be used to enhance differentiation of cardiomyocytes from myoblast progenitor cells.

We use the H9c2 cell line as a model system for cell differentiation[37]. Originally derived from rat embryonic ventricular heart tissue, this spontaneously immortalized line differentiates towards skeletal muscle upon reaching confluence[37]. Yet, upon exposure to retinoic acid, cardiac differentiation can be recovered[38]. The cardiac differentiation efficiency of H9c2 cells can be further enhanced by the presence of fibroblast-derived matrix in addition to the retinoic acid[39]. Our goal is to optimize cardiogenic differentiation of H9c2 by combining fibroblast support with appropriate mechanical and matrix cues in composites of coagulated fibrin and porcine cardiac dECM hydrogels. We aim at replacing the exogenous retinoic acid by endogenous cues to simplify the differentiation protocol. This allows to avoid the presence of the strong, widespread effects of retinoids on gene expression in screening and gene expression profiling experiments[40].

Regarding the development of a cardiac *in-vitro* niche with relevant cellular, mechanical and matrix components, a major aim is also to better support physiological electromechanical activity and synchronization of primary neonatal cardiomyocytes. Hence, we investigate synchrony, beating rate and recovery time of neonatal cardiomyocytes in various hydrogel compositions including fibrin, collagen I and matrigel, but also the composites fibrin-collagen I and dECM-fibrin. This allows us to further refine our comprehension of niche effects, with relevance to culture of primary neonatal cardiomyocytes and the optimal definition of tissue engineering and transplantation matrices.

## 2. Materials and methods

### 2.1. Extracellular matrix decellularization and characterization

Decellularized porcine cardiac extracellular matrix forms the basis of the dECM-fibrin hydrogel studied in this report. The procedure for decellularization of the cardiac tissue was based on published literature[10]. Briefly, porcine heart tissue was obtained from a local slaughterhouse and the ventricles cut into pieces of about 1 mm in thickness. The pieces were rinsed with deionized water and then stirred in 1% Sodium Dodecyl Sulfate (SDS) in a phosphate buffered saline (PBS) solution for 48-72 h at 4°C, followed by 1% Triton X-100 for an additional 30min. Finally, the preparation was stirred in deionized water overnight and freeze-dried. The dECM powder thus obtained was suspended in 0.1M HCl, followed by pepsin (Sigma P6887) digestion for 72 hours (100mg of dECM and 10mg of pepsin per 1mL of HCl)[10]. The pH was then adjusted to 7.4 by gradual addition of NaOH, yielding a solution of 100 mg/mL dECM. dECM stock solution was finally obtained by adding Dulbecco’s Modified Eagle Medium DMEM (Thermofisher, cat# 41965) to reach a final dECM concentration of 50mg/mL.

To verify the extent of decellularization, the residual DNA content in both native and decellularized tissue was measured. For this, dECM (respectively intact cardiac tissue) was dissolved in lysis buffer (0.5 M EDTA pH 8.0, 0.5% SDS) and 100μg/mL proteinase K (Sigma, P4850) overnight at 55◻[41]. The resulting suspension was vortexed, and proteins precipitated with phenol-chloroform, followed by centrifugation at 13700rpm for 40min. DNA was then isolated by recovery of the top (aqueous) phase, followed by addition of 0.5 mL ethanol, resuspension in deionized water, and quantification by Carry 50 spectrophotometer using a quartz cuvette at 260nm. For further quantification of dECM composition, collagen and Glycosaminoglycan (GAG) content in the dECM were measured using the Sircol™ Soluble Collagen Assay kit and Blyscan Sulfated Glycosaminoglycan Assay kit. Finally, histological sections of dECM were obtained by standard paraffin embedding. The sections were stained with hematoxylin-eosin (H&E), Sirius red, and Miller staining and scanned using an Olympus VS120-L100 microscope slide scanner to verify the presence of collagen and elastin.

### 2.2. Hydrogel preparation

To produce the compound dECM-fibrin hydrogel, we first prepare a pre-gel solution. For this, we mix 100 μl of 50mg/mL dECM stock solution (as described above), 528 μl of 50mg/mL fibrinogen (Sigma, F3879) solution, 100μl of HEPES (4-(2-hydroxyethyl)-1-piperazineethanesulfonic acid) 1M pH 7.4 and 269 μl of DMEM. To induce gelling of the pre-gel solution, we then add 1.7 μl of Thrombin (Sigma, T1063, 250U/mL) and 1.3 μl of calcium chloride 1M, and start incubation at 37 ◻. This yields a final composite gel with 5mg/mL dECM and 26.4 mg/mL fibrinogen.

Matrigel (Sigma, E1270, solution supplied at 9mg/mL) was diluted to 3mg/mL before gelation by using DMEM. Collagen I (Sigma, C4243, solution supplied at 3mg/mL) was diluted to 1mg/mL with DMEM prior to gelation. This also neutralized the pH. Finally, the fibrin-collagen I composite was prepared by mixing 157 μl of 3mg/mL collagen I stock solution, 314 μl of DMEM, followed by 528 μl of 50mg/mL fibrinogen. Gelling was then induced by adding 1.3 μl of calcium chloride and 1.7 microliters of thrombin (250U/mL).

To include cells, the necessary amount of cells to achieve a final total cell concentration of 10^6^ cells/mL was pelleted, followed by complete removal of the supernatant. The pellet was resuspended in pre-gel, followed by induction of gelation by addition of calcium chloride and thrombin solutions (fibrin-based hydrogels), and placement at 37◻ in 5% CO_2_ atmosphere.

### 2.3. Mechanical properties

To measure the mechanical properties, fibrin and dECM-fibrin hydrogels were prepared and pipetted into cryovials where they remained for 30 min at 37°C to gel. The samples were then removed from the cryovials and subjected to compression testing using a dynamic mechanical analyzer (TA Instruments DMA Q800). The storage and loss moduli are defined as follows[42]:

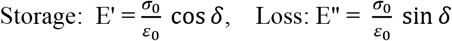

Where σ_0_ is stress, ε_0_ is strain, and δ is the phase angle or phase lag between the stress and strain. The Young’s modulus is evaluated as: 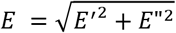

### 2.4. Microstructure characterization

The microstructure was analyzed by scanning electron microscopy (SEM). For this, the hydrogel sample was frozen and lyophilized. The lyophilized sample was coated with a few nanometers (5-10nm) of gold prior to performing SEM imaging.

### 2.5. dECM stability study with fluorescently labeled dECM

To investigate the stability of the dECM in our composite hydrogel, we fluorescently labeled the dECM using rhodamine isothiocyanate[43]. For this, we suspended 300mg dECM powder in 10mL of 0.1M HCl. The pH was adjusted to 10.3 with 0.9mL NaOH 1M and 0.3mL of Na_2_CO_3_ 1M. 6mg of rhodamine isothiocyanate was dissolved in 10mL of isopropanol, and this solution was added to the reaction mix, followed by 10mL of distilled water. After overnight incubation, the dECM was precipated and repeatedly washed with excess isopropanol, until a clear washing solution above strongly stained precipitate was obtained. The precipitate was air-dried overnight. dECM-fibrin hydrogels were prepared with fluorescently labeled dECM, using identical procedures as for unlabeled dECM. Fluorescent imaging was conducted after 1, 7, and 14 days of incubation in PBS at 37°C.

### 2.6. *In-vitro* cell studies

#### 2.6.1. H9c2 cell culture

H9c2 cells were obtained from the European Collection of Authenticated Cell Cultures (ECACC) (Lot# 17A028). The cells were cultured in DMEM medium supplemented with 10% fetal bovine serum and 1% penicillin and streptomycin in 75 cm^2^ tissue culture flasks at 37°C and 5% CO_2_ in an incubator. In accordance with supplier instructions, the H9c2 cells were passaged before reaching 70-80% confluency to avoid loss of differentiation potential[38].

#### 2.6.2. Nor-10 cell culture

NOR-10 (ECACC 90112701) cells were obtained from the European Collection of Authenticated Cell Cultures. The cells were cultured in DMEM medium supplemented with 10% fetal bovine serum and 1% penicillin and streptomycin in 75 cm^2^ tissue culture flasks at 37°C and 5% CO_2_ in an incubator. They were split before reaching 70-80% confluency according to supplier’s notice[44].

#### 2.6.3. H9c2 differentiation

As a starting point in our investigation into use of various hydrogels to enhance cardiogenic differentiation of H9c2 cells, we used a known differentiation procedure of H9c2 cells to cardiomyocytes[45]. This procedure implies simultaneous decrease of the serum percentage to 1% and application of retinoic acid for 5 days, before one to two last days of culture in expansion medium (DMEM with 10% FBS without addition of retinoic acid)[45]. Using this protocol, we studied co-culture with Nor-10 fibroblasts (100/0, 70/30, 50/50, 30/70, 0/100 H9c2:Nor-10 ratios with the total cell density of 10^6^cells/mL) in different concentrations of retinoic acid (0-2000nM) for enhancement of the cardiogenic differentiation efficiency. With a fixed 30% H9c2, 70% Nor-10 ratio and in the absence of retinoic acid, we then screened various hydrogels such as collagen I, matrigel, and the thrombin-coagulated dECM-fibrin hydrogel for their capacity to substitute for retinoic acid in H9c2 cardiogenic differentiation.

#### 2.6.4. Cell seeding onto hydrogel surfaces (2D)

To study the biochemical influence of different hydrogels on cardiomyocyte differentiation in a 2D geometry, we prepared ca. 0.5mm high hydrogel blocks in 48-well plate. For this, we gelled 50μl of dECM-fibrin, collagen I or matrigel in wells of interest. We then applied H9c2 cells mixed with Nor-10 fibroblasts at a ratio of 30% to 70% at a density of 2.5×10^5^ cells per cm^2^ on top of the hydrogels. The differentiation protocol was conducted without addition of retinoic acid: 5 days with 1% FBS in DMEM, followed by 1-2 days of 10% FBS in DMEM[45].

#### 2.6.5. Cell seeding in 3D hydrogel

To assess whether differentiation could be improved in 3D vs. 2D, we mixed dECM-fibrin pre-gel with Nor-10 and H9c2 cells (70/30 ratio, 10^6^ cells/mL). Then thrombin and calcium chloride were added to the solution, and 200μl of the solution was rapidly poured in a 48-well plate where the hydrogel solidified in the incubator. Collagen I and Matrigel were used as controls. The differentiation protocol was otherwise identical to the differentiation experiments on the 2D hydrogels, with serum reduction only, in the absence of exogenous retinoic acid.

#### 2.6.6. Immunostaining and 3D imaging

For immunostaining, cell culture samples were fixed in 4% of paraformaldehyde (PFA) for 20min at room temperature. Then, 0.1% TritonX-100 was added to permeabilize the cells for 30min at room temperature. By incubating cells with phalloidin-Atto 488 (Sigma 49409, 1:50) for 45min at 4°C, actin filaments were made visible in all types of cells [46].

We assessed the differentiation by measuring the fraction of H9c2 cells positive for troponin T by immunofluorescence using the T6277 antibody from Sigma. As this antibody detects both cardiac and skeletal muscle troponin T, we also confirmed cardiac differentiation on dECM-fibrin samples and tissue culture plates using an antibody specific to cardiac troponin T (Abcam, ab8295). This data is provided in supplementary figure S1.

For immunofluorescence, the cells were first blocked with 1% BSA for 1 hour at 37°C. Troponin T primary antibody (Sigma, T6277, 1:50) was then added, followed by incubation overnight at 4°C. Alexa-568 secondary antibody (Sigma, A10037, 1:1000) was added after washing with DPBS (Gibco 2062235) and incubated for 45min at 37°C, prior to washing with DPBS and staining with 4’,6-diamidino-2-phenylindole (DAPI, 1:2000 from 5mg/ml stock solution) for 5min. DAPI was replaced with DPBS before imaging the cells under a fluorescence microscope. We quantify the differentiation percentage as the surface area covered by cells expressing troponin T, relative to the total area of the fluorescent images, and normalize to the percentage of H9c2 cells seeded in co-culture.

#### 2.6.7. Neonatal cardiomyocytes isolation

To study the effect of the thrombin-coagulated dECM-fibrin hydrogel on primary cardiomyocytes, we isolated primary rat neonatal cardiomyocytes using the Pierce Primary Cardiomyocyte Isolation Kit, according to the manufacturer’s instructions[47]. Organ harvesting was performed on donated sacrificed control animals from unrelated experiments, according to license VD 3290 designed for this specific purpose.

#### 2.6.8. Calcium imaging

For calcium imaging, we seeded the primary rat cardiomyocytes (2.5×10^5^cells/cm^2^) onto hydrogel slabs prepared in 48-well plates (50 μl of hydrogel per well, as before). Tissue culture controls were prepared by leaving out the hydrogel polymerization step. After 3 days of culture in DMEM with 10% FBS, the cultures were loaded with 2 μM Fluo-4 AM (Sigma, F14217) for 30 min at 37°C. Calcium transients were then recorded using fluorescent microscopy at room temperature. Temporal peak detection was based on a custom ImageJ plugin, implementing the publicly available findpeaks function of Octave[48] in Java. The plugin is available as supplementary S2, but also for download at https://github.com/tbgitoo/calciumImaging, along with source code and a user manual. We used this plugin to evaluate local beating frequency and temporal phase shift from the calcium imaging videos.

#### 2.6.9. Beating characteristics

To assess the contractile properties of primary cardiomyocytes in 3D cultures, we suspended the primary rat cardiomyocytes at a cell density of 10^6^ cells/mL in pre-gel mixtures of dECM-fibrin, fibrin, and fibrin-collagen I composite. After gelling of 200 μL of cell-hydrogel mixture per well in a 48-well plate, the cultures were followed both visually and videographically. Synchrony and onset of beating was judged visually on a per-well basis. To quantify the mechanical beating rate of the cells, movies were acquired by connecting a video camera to the microscope, while maintaining the samples at 37℃ in a temperature-controlled chamber. For quantification of the beating frequency, we used the Pulse software (Cellogy Inc.)[49].

### 2.7. Statistical analysis

Data were compared using unpaired t-test (two-tailed, equal variances) in the GraphPad software. Error bars represent the mean ± standard deviation (SD) of the measurements (∗ p < 0.05, ∗ ∗ p < 0.01, and ∗ ∗ ∗ p < 0.001).

## 3. Results

### 3.1. Development and characterization of the hybrid hydrogel

#### 3.1.1. Decellularization characteristics

We successfully decullarized porcine cardiac extracellular matrix. As shown in Fig. 1A, the total amount of remaining DNA is less than 50ng/mg of tissue, indicating essentially complete removal of cells[50]. The concentration of the extracellular matrix components collagen and glycosaminoglycans (GAG) was measured in he native tissue and after decellularization, and shows no loss of these components in the resulting dECM (Fig. 1B, 1C). H&E, Sirius red for collagen and Miller for elastin staining confirmed the absence of cells and cell debris and the presence of collagen and elastin in the matrix after decellularization (Fig. 1D).

**Figure 1.**
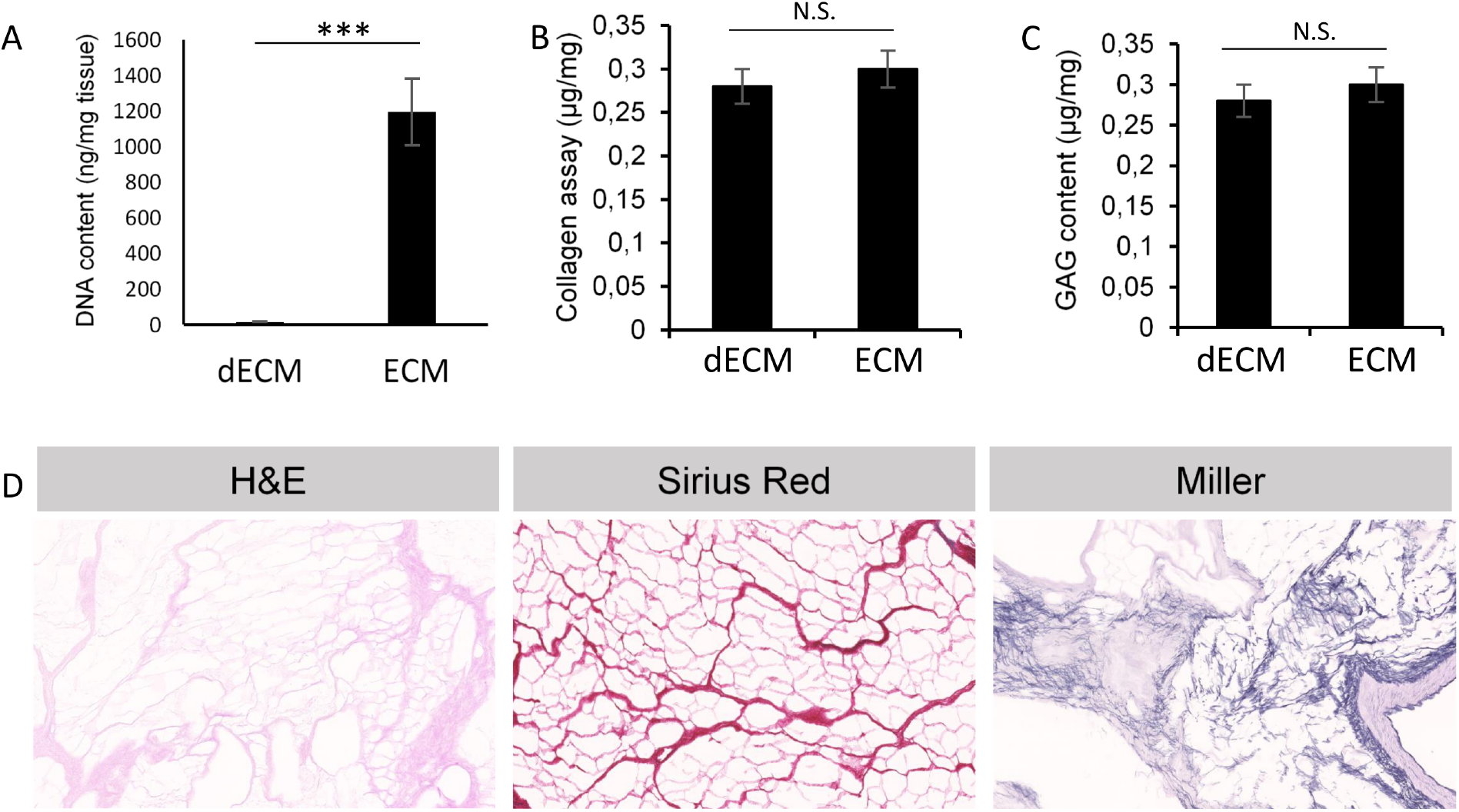
dECM characterization. A) DNA, B) collagen, and C) Glycosaminoglycane (GAG) content of the dECM. D) H&E, Sirius red and Miller staining used for staining the nuclei, collagen and elastin respectively, which proves the presence of collagen and elastin and removal of the DNA. *** = significant difference for p < 0.001 (N = 3-4 per condition). Scale bar: 100μm.

#### 3.1.2. Mechanical properties

Fig. 2A shows the relation between the elastic modulus and fibrinogen concentration in pure fibrin gels. In agreement with literature[12], higher fibrinogen concentrations are associated with higher Young moduli, although at the highest concentrations, a saturation effect can be seen. The evaluation of the storage and loss moduli in dECM-fibrin gels in a linear temperature scan is presented in Fig. 2B.

**Figure 2.**
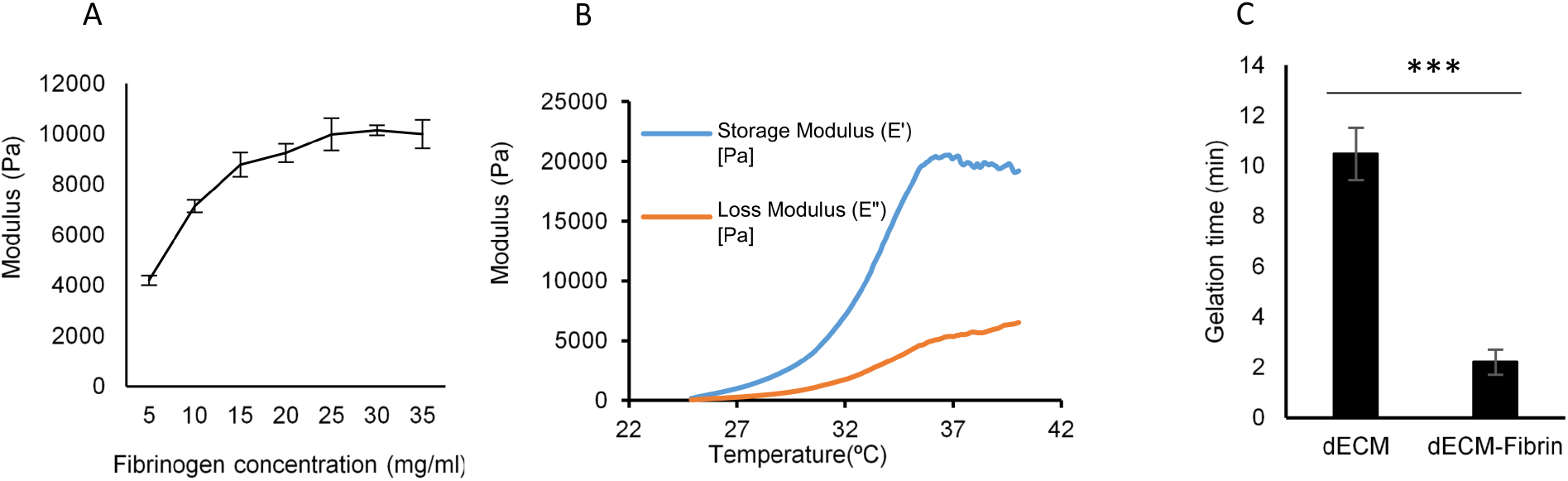
Mechanical characterization. A) Stiffness of fibrin gels at different concentrations at room temperature. This allows to determine the optimum concentration of fibrinogen in the hydrogel. B) Storage and loss moduli of the dECM-fibrin hydrogel in a temperature scan (from 25 to 40). The Young’s modulus of the hydrogel can be calculated from these measurements. It is comparable to the one of the native heart. C) Gelation time of the dECM and dECM-Fibrin hydrogels at 37◻. In the case of the dECM-Fibrin hydrogel, 2 minutes are sufficient for handling the gel *** = significant difference, p < 0.001 (N = 3-4 per condition).

At 37◻, the Young’s modulus of the dECM-fibrin hydrogel stabilizes at about 21 kPa, falling into the reported range of native adult myocardium from 11.9 to 46.2 kPa[8]. This confirms that mechanical properties adequate for heart cell culture can be achieved with the thrombin coagulation method chosen, eliminating the need for less specific crosslinking agents[8], [51].

#### 3.1.3. Gelation time

Fig. 2C shows that the gelation time of dECM alone, in the absence of cells, is about 10 minutes. By adding fibrinogen and thrombin to the hydrogel, the gelation time is reduced to about 1-2 minutes, which is sufficient for handling the cells in 3D while avoiding cell sedimentation[27].

#### 3.1.4. Structural characterization and dECM stability

Fig. 3A shows a scanning electron micrography image of the dECM-Fibrin hydrogel, whereas Fig. 3B shows specifically the spatial distribution of rhodamine-labelled dECM within the dECM-fibrin composite. The heterogeneous structure of the hydrogel can be observed in both imaging modes. This implies that coherent pieces of dECM subsist and are embedded into fibrin, rather than forming a spatially homogeneous mixture, with heterogeneity probably defining local niche structures. Fig. 3C and 3D indicate stable maintenance of dECM within the dECM-fibrin composite at 7 and 14 days, with no loss of dECM detected. The confocal stack of control dECM-fibrin hydrogel without rhodamine labeling shown in Fig. 3E validates the specific detection of rhodamine-labelled dECM as it shows virtual absence of autofluorescence at the exposure settings used for Fig. 3B-3D.

**Figure 3.**
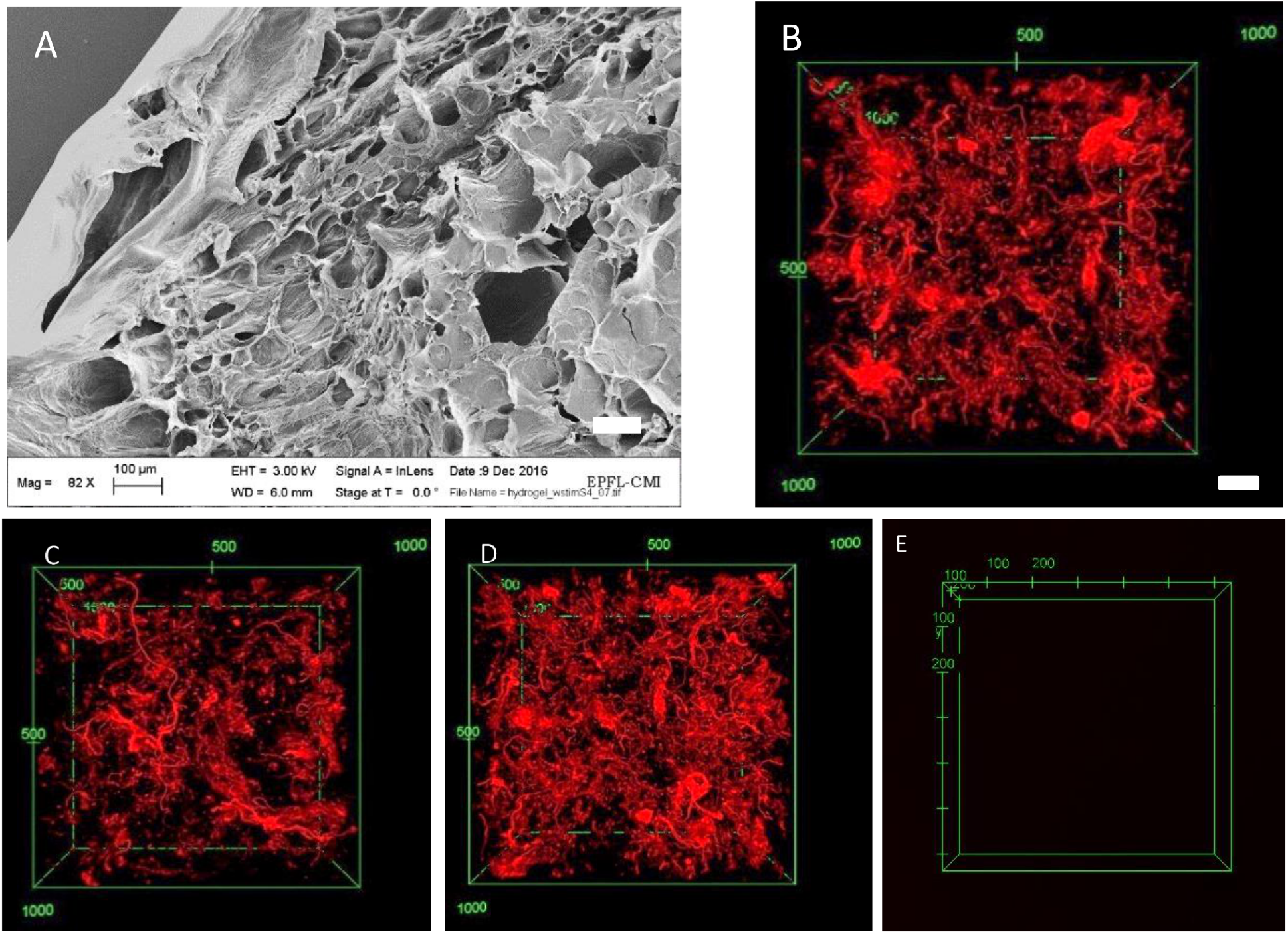
Microstructural analysis of dECM-fibrin gels. A) Scanning electron microscopy image of the dECM-fibrin hydrogel. B-D) dECM was labeled with rhodamine isothiocyanate and gelled together with fibrin. Confocal images were taken after B) 1 day, C) 7 days, and D) 14 days, E) the dECM-fibrin hydrogel without rhodamine-labelling. The distribution of dECM is locally heterogeneous. No significant loss of dECM occurs during this time. Scale bar: 100μm.

### 3.2. H9c2 in co-culture with fibroblast cells

To investigate the role of extracellular matrix components, and particularly dECM-fibrin as compared to control matrices such as collagen and matrigel, we used cardiogenic differentiation of the H9c2 myoblast line as a model system[52]. The differentiation of H9c2 cells to cardiomyocytes is traditionally initiated by reducing the serum in the culture medium to 1% in the presence of all-trans-retinoic acid[45].

In preliminary trials to adapt this procedure for use with our hydrogel systems, we found that retinoic acid induces fibrinolysis. This led to dissolution of fibrin-based gels. Retinoic acid is a known transcriptional activator for the expression of fibrinolytic enzymes[53][54]. In addition, we find the dissolution effect to partly persist even in cell-free systems, indicating a chemical effect as well. Hence, our first step is to optimize cardiac differentiation in H9c2 cells in the absence of retinoids.

Fibroblast-derived matrix has been shown to enhance retinoid-based differentiation of H9c2 cells[39]. Here, we assess whether a similar favorable effect on the H9c2 cardiogenic differentiation can be obtained by direct co-culture with Nor-10 fibroblasts[55].

We first optimize the co-culture ratio of H9c2 cells to Nor-10 fibroblasts. The aim is to increase cardiogenic differentiation of H9c2 cells in the presence of a minimal amount of retinoic acid. For the sake of simplicity, we investigate this in 2D co-culture, without hydrogel, but systematically vary both the H9c2:Nor-10 seeding ratio and retinoic acid concentrations.

Table 1 shows the percentage of cardiogenically differentiated H9c2 cells, evaluated as the relative surface area covered by cells expressing troponin T corrected for the proportion of H9c2 seeded. Confirmation of specific cardiac troponin T expression is further given in Supplementary Figure S1B. The heatmap (Table 1A), outlines the differentiation percentage of the H9c2 cells thus calculated for different values of the ratio of fibroblast cells to H9c2 cells and different concentrations of retinoic acid after 7 days in culture. The results demonstrate that the lower the fraction of H9c2 seeded, the higher the relative differentiation efficiency, indicating that neighboring Nor10 cells help drive the cardiac differentiation of H9c2 cells. As a compromise between high relative differentiation efficiency and achievable absolute numbers of differentiated cells, we choose the seeding ratio of 30% H9c2 cells to 70% fibroblast cells as the preferred condition for subsequent experiments. Of note, even this optimized co-culture could not completely substitute for the addition of retinoic acid, as below 62.5nM the differentiation efficiency started dropping, to reach only about 7% in the absence of the retinoid (Table 1B). Overall, we retain that performing co-culture with fibroblasts improves the differentiation efficiency and dramatically reduces the required concentration of retinoic acid for differentiation.

**Table 1.**
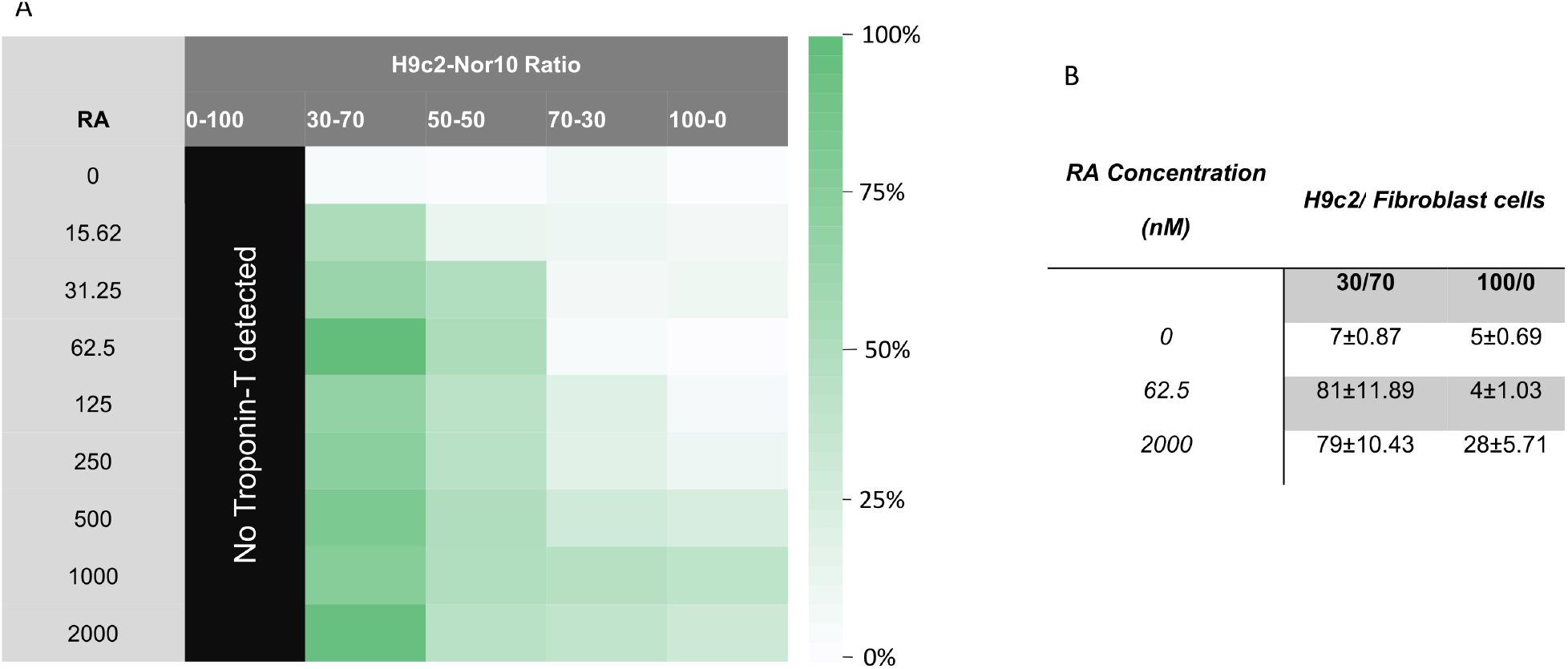
Differentiation of H9c2 cells in mono-and co-culture on tissue culture plates. A) The heat map of normalized differentiation percentage of H9c2 cells for different concentrations of retinoic acid and for different ratios of H9c2 to fibroblast cells. Cultures in 2D, without hydrogel, for 7 days (differentiation duration). B) Selected numerical values for the differentiation percentages for the 30/70 ratio and 100/0 ratio.

### 3.3. Cell seeding and differentiation in 2D on hydrogel surfaces

We next assessed the capacity of various hydrogels to enhance cardiogenic differentiation in H9c2 cells. We also aimed at replacing the retinoic acid in the H9c2 differentiation protocol by similar favorable effects mediated by the microenvironment.

Having confirmed that Nor-10 fibroblasts favor cardiogenic differentiation of H9c2 cells, we seeded H9c2 cells and Nor-10 fibroblasts at a ratio of 30%:70% onto hydrogel slabs in 48 well plates. Fig. 4A to 4D show the extent of troponin T expression on dECM-fibrin, matrigel, collagen and tissue culture plate control after 1 week of differentiation (confirmation for specific cardiac troponin T expression in Supplementary Figure S1C). Fig. 4E quantifies the percentage of differentiation of H9c2 in co-culture with fibroblasts on the different materials. The proportion of differentiated H9c2 cells is larger on the dECM-fibrin hydrogel than on cell culture plates (P=0.0001) and matrigel (P=0.0024), but similar to the one on collagen (P=0.645). These results suggest that the influence of dECM on the differentiation of H9c2 cells is likely mediated by its collagen content.

**Figure 4.**
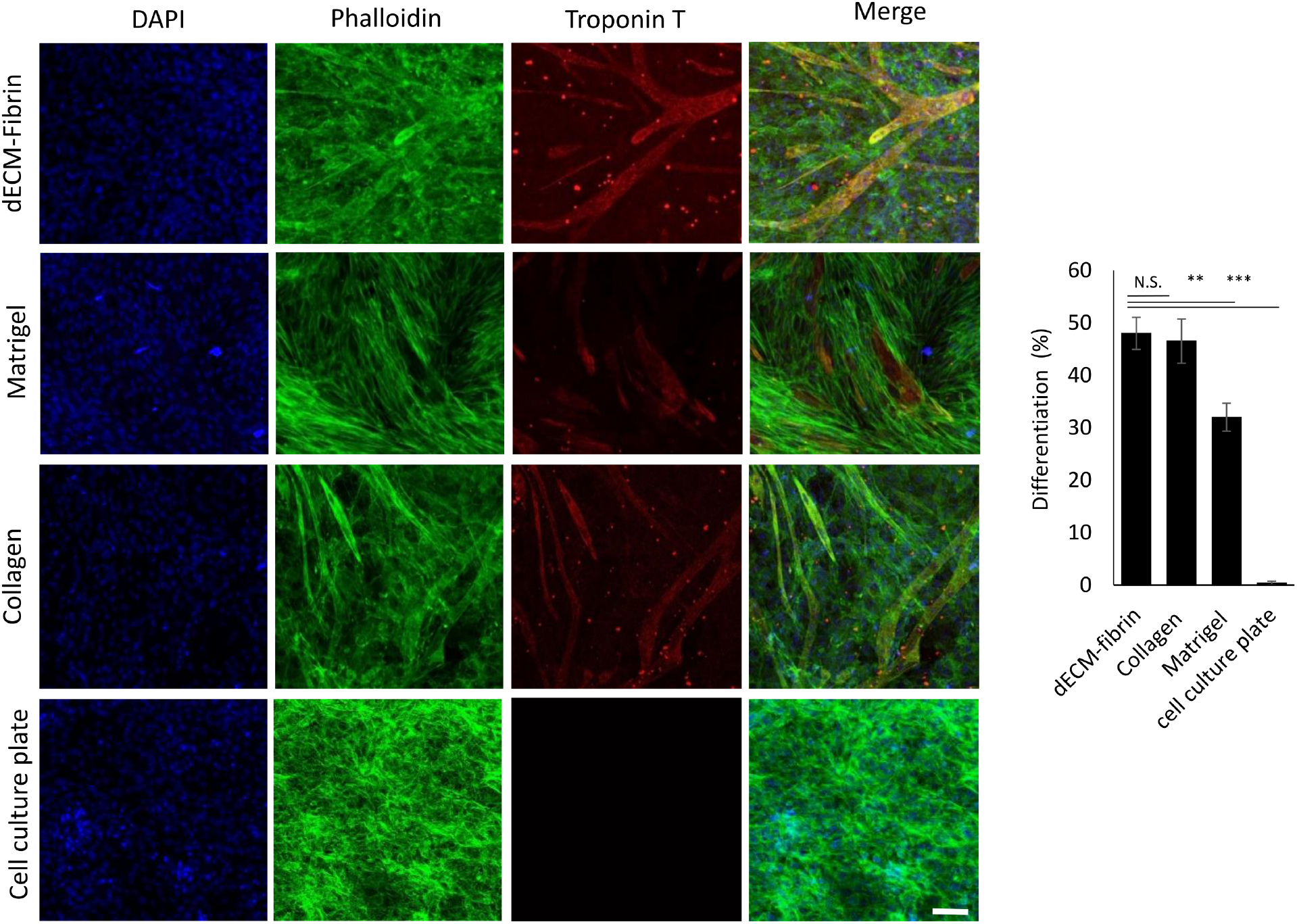
Cifferentiation of H9c2 cells in co-culture with fibroblasts on different hydrogels in the absence of retinoic acid. Differentiation A) on dECM-fibrin, B) on Matrigel, C) on collagen I, and D) on cell culture plates after 7 days of differentiation E) Percentage of differentiation of H9c2 cells on different hydrogels. Troponin T staining in red shows the differentiated cells, phalloidin stains the actin filaments of both fibroblasts and H9c2 cells in green. DAPI stains the nuclei of the cells. Scale bar: 100μm (P=0.0024 for dECM-fibrin vs. Matrigel, P=0.0001 for dECM-fibrin vs. cell culture plate,, and P=0.645 for dECM-fibrin vs. collagen).

The effect of suitable extracellular matrix is striking: The combination of co-culturing the H9c2 cells with Nor-10 fibroblasts and the presence of either dECM-fibrin or collagen outperforms even the highest concentrations of retinoic acid in H9c2 monoculture in 2D (comparison to Table 1B, 100/0 ratio, 2000nM retinoic acid, P=0.0059 between dECM-fibrin and 2D, and P= 0.0106 between collagen and 2D).

### 3.4. Cell seeding and differentiation in 3D hydrogels

Having evaluated and optimized differentiation efficiency on the surface of various hydrogels, our next step was to proceed to a truly 3D co-culture configuration. For this, we included the suspended cells H9c2 and Nor-10 cells to be co-cultured in the dECM-fibrin hydrogel prior to gel formation. Cells cultured in the 3D structure of the dECM-fibrin hydrogel attached, spread, and formed a network throughout the wells. We also attempted 3D culture in collagen I and matrigel[56], [57], but could not carry out the entire differentiation protocol in 3D, since after about 2 days, the gels had become very soft with a large part of the cells attached to the floor of the wells.

Fig. 5 shows the H9c2 cell differentiation in the 3D hydrogel using Troponin T staining. The cells attached, spread, and formed a network throughout the wells. The differentiation percentage of H9c2 cells in co-culture with fibroblasts in 3D dECM-fibrin hydrogel was 83±8%. This is significantly better than the 2D co-cultures of H9c2 and fibroblast cells on top of dECM-fibrin coated surfaces (P = 0.0007), indicating a specific advantage of the 3D configuration. To achieve a similar differentiation efficiency in 2D, the combination of retinoic acid and co-culture is required (P=0.77).

**Figure 5.**
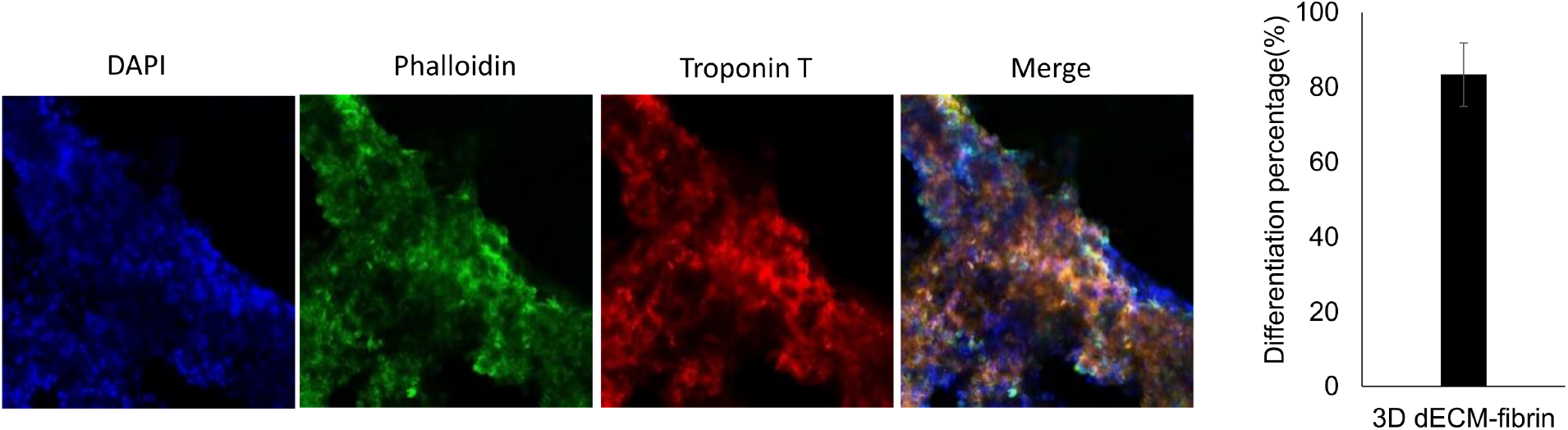
Co-culture of 30% of H9c2 cells and 70% fibroblasts in a 3D dECM-fibrin hydrogel. A) Confocal images of the differentiation of H9c2 cells in the dECM-fibrin hydrogel after 7 days (differentiation duration). Troponin T staining shows the differentiated cells in red. Phalloidin stains the actin filaments of all the cells in green and DAPI in blue stains the nuclei. Scale bar: 100μm. B) Percentage of differentiation of the H9c2 cells in the co-culture with fibroblast cells in the 3D dECM-fibrin hydrogel after 7 days (differentiation duration).

Due to the fragile nature of the pure collagen I gels, we could not assess whether there is a difference between dECM-fibrin and collagen I regarding H9c2 differentiation in 3D. However, from the 2D results given in Fig. 4E, it seems that the primary advantage of the dECM-fibrin gel over collagen I in the H9c2 system is mechanical ruggedness, both gels showing high differentiation capacity.

Overall, we conclude that the very high differentiation efficiency in our 3D co-culture system affords the freedom to avoid retinoids altogether. Co-culture with Nor-10 fibroblasts, the 3D environment, and a collagen-based matrix all have a positive effect on cardiac differentiation of H9c2 cells.

### 3.5. Neonatal cardiac cells on 2D and in 3D hydrogels

To study the effect of different hydrogels with a more physiologically relevant cell system, we next cultured primary neonatal cardiomyocytes on surfaces (2D culture) of dECM-fibrin hydrogel, in comparison with various controls: tissue culture plates, the commercial matrices collagen I and matrigel, as well as a fibrin-collagen I composite. This latter is the closest analog to dECM-fibrin with chemically defined composition. To assess electrophysiological activity, we videographically recorded intracellular calcium ionic concentration variations by using the Ca^2+^ indicator Fluo-4 AM at 3 days of culture. Sample videos are provided as supplementary Videos S4 (dECM-fibrin), S5 (fibrin-collagen I), S6 (collagen I), S7 (Matrigel), S8 (tissue culture plate). From the calcium oscillation videos, we evaluated local frequency (Fig. 6A-6E) and local phase (Fig. 6F-6J). The results indicate essentially perfect synchrony on the dECM-fibrin hydrogel, regarding both frequency (6E) and phase (6J). This is followed by fibrin-collagen I where the phase analysis (6I) shows some individual desynchronized cells and regions with increased lag. The other conditions show even higher degrees of variability in phase and frequency, common timing being nearly lost on matrigel (6G). This indicates that dECM-fibrin performs best regarding synchronization of calcium influx. The result correlates with morphological observation: cell spread and morphological connectivity improve in parallel with synchrony along the series matrigel, collagen I and tissue culture plates, fibrin-collagen I, culminating in dECM-fibrin hydrogels.

**Figure 6.**
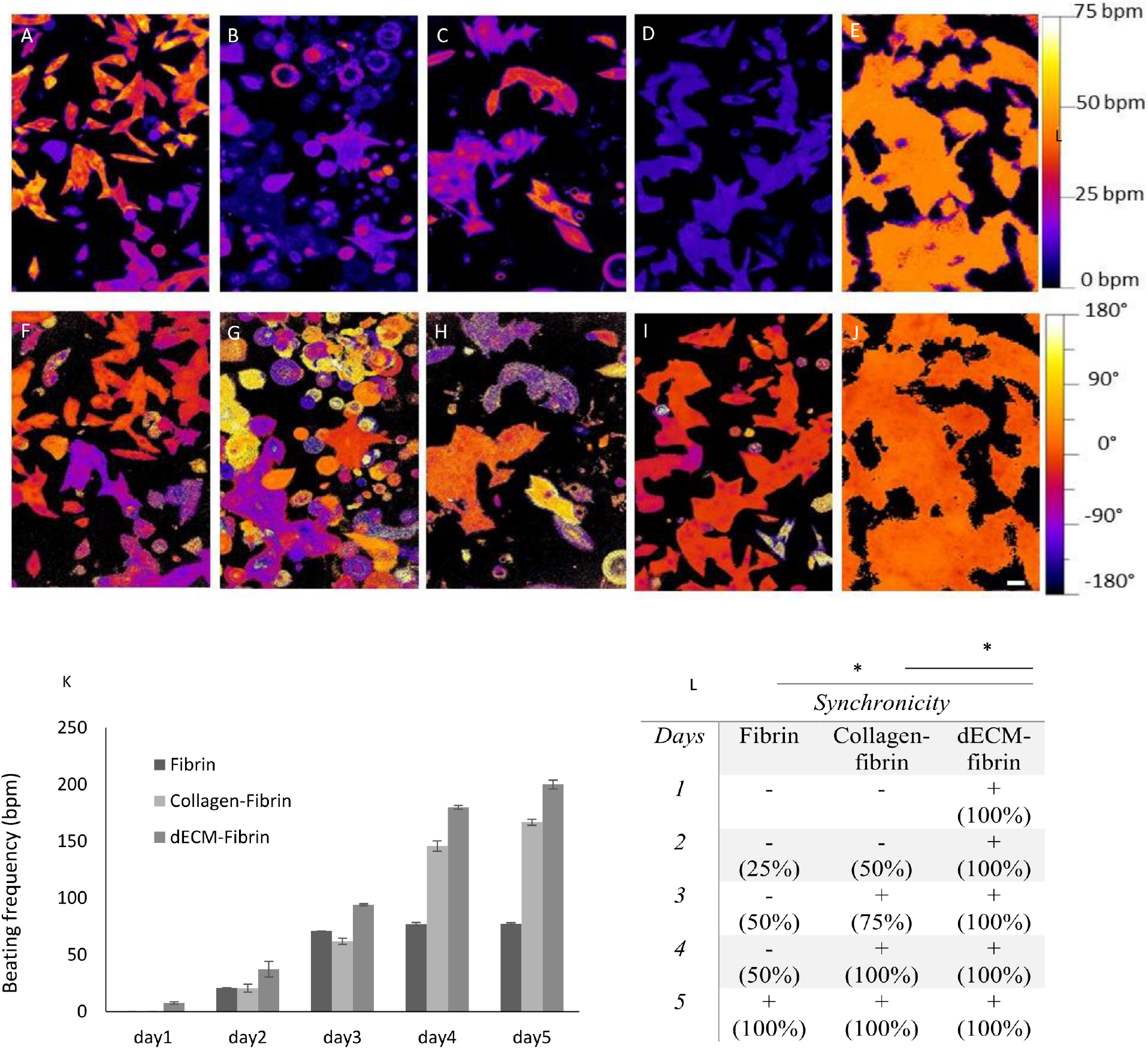
Calcium transients and beating characteristics of cardiomyocytes interacting with different hydrogels. A-J) Calcium imaging for neonatal cardiomyocytes seeding onto different hydrogels, and tissue culture plate (control). A-E) Frequency, F-J) Phase (positive values indicate earlier beating, negative value retardation). Hydrogels used in calcium imaging: tissue culture control (A and F), matrigel (B and G), collagen I (C and H), fibrin-collagen I (D and I) and dECM-fibrin (E and J). K) Beating rate for the first 5 days for neonatal cardiomyocytes seeded 3D in fibrin, fibrin-collagen I and dECM-fibrin hydrogels. L) Synchronization as the percentage of synchronously beating wells (four wells per condition) during 5 days in the 3D hydrogels. Scale bar: 100μm.

Having established optimal properties of the dECM-fibrin matrix regarding synchronization of calcium influx, we next investigated mechanical contraction activity. In light of the results obtained by calcium imaging, we compared 3D cultures of rat neonatal cardiomyocytes in dECM-fibrin hydrogels to similar cultures in fibrin alone and fibrin-collagen I.

The normal beating rate of the neonatal rat heart is around 276-423 beats per minute (bpm)[58], whereas for isolated primary cells *in-vitro* on standard cell culture conditions it is about 100-115 bpm, (manufacturer’s notice[59], confirmed in preliminary trials). Fig. 6K shows a profound effect of the extracellular matrix environment on the beating rate. At 77 bpm at 5 days of culture, pure fibrin gels sustain a relatively low beating rate, whereas fibrin-collagen I (166bpm, P=0.0001 vs. fibrin) and even more so dECM-fibrin cultures (206bpm, P=0.0001 vs. fibrin-collagen I) approach the expected rate of the neonatal heart. The recovery of the mechanical beating function was gradual as indicated by the progressive rise of beating frequency for all conditions in Fig. 6K, and started earlier in the dECM-fibrin hydrogel as compared to the other conditions.

Finally, in Fig. 6L, we visually quantified the synchrony of the neonatal cardiac cell cultures. The neonatal cardiomyocyte extracts seeded in dECM-fibrin gel displayed synchronous contractions from the first day on, and for at least 10 days (Supplementary video S3). In contrast, cardiomyocytes seeded in fibrin did not show synchronous beating until day 5 and typically stopped beating after 7 days. The cardiomyocytes cultured in fibrin-collagen I showed intermediate behavior by reaching synchronicity at day 3, but similar to fibrin, they did not beat after 7 days. These results confirm the superior results obtained with the dECM-fibrin hydrogel in calcium imaging (Fig. 6A-6J), and make dECM-fibrin the optimal hydrogel for primary neonatal cardiomyocyte culture among the various options compared here.

## 4. Discussion

In this study, we develop a hydrogel blend of fibrin and decellularized porcine cardiac matrix to improve the cellular niche for cardiomyocyte cell culture. For this, we successfully decellularized porcine ventricular tissue, with maintenance of collagen, glycosaminoglycanes and elastin components. The addition of fibrinogen to the dECM thus obtained, followed by coagulation with thrombin, provided a Young modulus of about 20kPa, in the range of adult cardiac tissue[8]. We used this system in conjunction with fibroblast co-culture to robustly differentiate the H9c2 myoblast line into a cardiomyocyte phenotype while avoiding retinoic acid. Finally, we demonstrated excellent maintenance of electromechanical activity in neonatal cardiomyocytes by quantification of beating activity as well as calcium imaging.

An important aim in the design of the hydrogel system was to match the physical stiffness of native myocardium (10-50kPa range, [60][8]). To mechanically reinforce the intrinsically soft cardiac dECM, we chose the specific coagulation of fibrinogen by thrombin, as opposed to more generic agents such as transglutaminase[30][61] or genipin[39][62] We find the thrombin-coagulated fibrin gels to be softer than similar gels crosslinked by transglutaminase[30], but we could compensate for this by increasing the fibrinogen concentration correspondingly. The specificity of thrombin[63] is advantageous for both *in-vitro* and anticipated *in-vivo* applications, and probably allowed us to avoid cellular toxicity observed in transglutaminase crosslinking[30].

We used the H9c2 cell line to investigate cardiogenic differentiation. By combining 3D embedding into dECM fibrin hydrogels with co-culture with Nor-10 fibroblasts, we obtained over 80% differentiation efficiency. While the utility of adding fibroblasts or fibroblast-derived matrix in H9c2 differentiation has repeatedly been demonstrated[64][29][60], the differentiation efficiency in our system is rather high compared to previous reports[64][39]. *Nota bene*, we obtain our results in the absence of retinoids. We achieved this by stepwise optimization of three key elements of the cardiac cell environment: choice of the extracellular matrix, its physical properties, and the co-culture composition. The result should simplify pharmacological screening, as retinoids are known to broadly affect cellular processes beyond specific cardiac differentiation[66]. Further, avoiding the strong transcriptional effects of retinoids[66] could also simplify investigation of communication between cardiac fibroblasts and cardiomyocyte, which today remains only partially understood[67].

The ultimate aim of an *in-vitro* cardiac model should be to replicate the electromechanical beating function. Comparative culture of freshly isolated rat cardiomyocytes on dECM-fibrin, fibrin-collagen I and fibrin control hydrogels indicates specific advantages of the dECM-fibrin composite as it performs best regarding recovery of physiological beating frequency and reacquisition of synchrony. In line with these results, dECM-fibrin also outperformed collagen I, matrigel, tissue culture plates, and also fibrin-collagen I regarding synchronization in calcium imaging. The result correlates well with morphological analysis: cell spreading and enhanced geometric contacts are overall associated with higher frequency and better synchrony. Given that we seed a native cardiomyocyte population including various pacemaker cells, these results replicate native cardiac physiology[68]: better connectivity directly provides better synchrony, but also better spread of the highest pacemaker frequency. This subtly graded response of the native cardiomyocyte population contrasts with the differentiation of the H9c2 cell line, where the presence of collagen was as efficient as the dECM-fibrin composite in inducing cardiogenic differentiation.

Interaction of cardiomyocytes with extracellular matrix proteins is known to be of prime importance for cardiac differentiation and electrophysiological maintenance. For example, a multitude of integrins with affinity for collagen I, but also laminin and fibronectin are expressed on cardiomyocytes[69][70][71][72]. It is fully conceivable that the presence of a more complete mixture of extracellular matrix components in the cardiac dECM is important not only for fine electrophysiological regulation, but also for cell spreading and thus geometrical connectivity of neonatal cardiomyocytes. In contrast, collagen I may suffice in H9c2 to induce a pro-cardiogenic, but ultimately non-functional cell state.

Mechanical aspects probably play a more important role in mechanically beating cells as well. Ideal energy transmission to the substrate is expected to occur at near physiological stiffness (neither too soft, preventing force development, nor too stiff, preventing substrate deformation)[73]–[75]. Mechanical feedback[71], could therefore help explain why the most physiological frequency and best synchrony for the neonatal cardiomyocytes are indeed found for the composite hydrogels, and not for the softer collagen, matrigel or pure fibrin or on the other hand the hard tissue culture plates.

## 5. Conclusion

In this work, we demonstrate a new combination of two natural hydrogels for myocardium regeneration: dECM from porcine ventricular heart tissue, admixed with fibrinogen and coagulated under cell-compatible conditions with thrombin. The mechanical properties are adjusted to match the ones of the native heart tissue, while the choice of thrombin as a specific crosslinking agent eliminates concerns of toxicity with more generic agents. We successfully test the system in the context of highly efficient, retinoid-free cardiac differentiation of the myoblast line H9c2. In neonatal cardiomyocytes, we show enhanced recovery, synchrony and beating frequency as compared to various controls, demonstrating specific advantageous of using the dECM-fibrin hydrogel over collagen analogs, fibrin alone, matrigel and tissue culture plates. We anticipate use of this system both as robust cardiac 3D cell model for drug screening, and as a building block for 3D printing, tissue engineering and transplantation in regenerative medicine.

## 6. Acknowledgements

The research presented in this paper is supported by the Swiss Government Excellence Scholarship ESKAS-Nr: 2015.1050, and the SNSF Professorship Grant PP00P2_163684. The authors thank Prof. Lashuel’s lab, especially Elena Gasparotto for providing neonatal hearts and Prof. Rohr’s lab, especially Regula Fluckiger Labrada for providing isolated neonatal cardiac cells, Dr. Marisa Jaconi for providing advice on cardiac cell biology and Fabien Bonini for scanning the histology slides. We also thank Arnaud Bertsch for his priceless support. Special thanks of gratitude go to Marina Braschler and Sara Ancel. Further, we acknowledge the services of the Bioimaging and Optics Platform (BIOP), Center of MicroNanoTechnology(CMi) and Interdisciplinary Centre for Electron Microscopy(CIME) at EPFL, and the Bioimaging Core Facility of the Faculty of Medicine of the University of Geneva.

## 7. Declaration of conflicts of interest

T. Braschler and P. Renaud declare financial interests in Volumina-Medical SA, Lausanne, Switzerland. The other authors declare no conflicts of interest.

## 8. Supplementaries

Supplementary files and raw research data are available on the Zenodo repository: https://zenodo.org/record/3556793#.XeAxJ797lR0

S1: Staining for cardiac Troponin T

S2: calciumImaging_.jar. Source code, documentation and user manual at https://github.com/tbgitoo/calciumImaging

S3: Beating motion of rat neonatal cardiomyocytes in dECM-fibrin

S4: calcium imaging: video of calcium oscillations in rat neonatal cardiomyocytes on dECM-fibrin

S5: calcium imaging: video of calcium oscillations in rat neonatal cardiomyocytes on fibrin-collagen I

S6: calcium imaging: video of calcium oscillations in rat neonatal cardiomyocytes on collagen I

S7: calcium imaging: video of calcium oscillations in rat neonatal cardiomyocytes on matrigel

S8: calcium imaging: video of calcium oscillations in rat neonatal cardiomyocytes on tissue culture plastic

## Notes

https://github.com/tbgitoo/calciumImaging

